# Quantitative partition of the rhizosphere microbiota assembly processes

**DOI:** 10.1101/2020.04.18.048595

**Authors:** Ning Ling, Chao Xue, Philippe Vandenkoornhuyse, Qirong Shen

**Affiliations:** Jiangsu Provincial Key Lab for Solid Organic Waste Utilization, Jiangsu Collaborative Innovation Center for Solid Organic Waste Resource Utilization, Nanjing Agricultural University, Nanjing 210095, Jiangsu, China; Université de Rennes 1, CNRS, UMR 6553 EcoBio, campus Beaulieu, Avenue du Général Leclerc, 35042 Rennes Cedex, France

## Abstract

The soil microbial reservoir and plant recruitment are predominant forces determining microbiota assembly in rhizosphere (i.e. active and passive processes respectively), but to date, no straightforward method to evaluate the respective contribution of forces to the rhizospheric microbiota assembly rules is available. We propose herein a promising way to quantitatively partition the assembling forces of rhizosphere microbiota using ordination metrics. We anticipate that this new method can not only weight the plants individual contributions to microbiota assembly in rhizosphere, but can also indirectly provide a way to quantitatively evaluate soil health by the contribution from plant selection.

---

The rhizosphere is of central importance not only for plant nutrition, health and quality but also for microorganism-driven ecosystem functioning and nutrient cycling in agroecosystems [1-3]. It is well established that plant species and soil type cooperatively shape the structure and function of microbial communities in the rhizosphere, indicating that both plant selection and the soil microbial reservoir confer forces that serve to structure the rhizosphere microbial community [4-7]. It is also well known that plants can recruit particular microorganisms to the rhizosphere from the soil reservoir through root exudates and signaling compounds as a deterministic process of rhizosphere assembly [8]. Measurements of the degree of plant influence on the assembly of the rhizosphere microbiome would offer an improved assessment of the co-evolved fraction of rhizosphere microorganisms. It has recently been demonstrated that plants can leave a ‘fingerprint’ of their endosphere microbiome on the soil reservoir [9]. In an environment where plant diversity is low, one would expect there to be a pronounced influence of plants on the soil reservoir. This would be reflected in a reduction of heterogeneity between microbial communities in the bulk soil, rhizosphere and the root-endosphere. A similar trend is expected when the soil microbial reservoir has been eroded, perhaps due to a biotic or abiotic stressor. In both these cases the apparent weight of the plant contribution to the rhizospheric microbiota is expected to vary (i.e. bulk soil, rhizosphere and endosphere microbiota more similar). Therefore, the measure of the weight of the respective plant filtering/recruitment coupled with the influence of the bulk soil microbial reservoir should allow the interpretation of the level of disturbance to the soil reservoir.

An experimental design where bulk soil, rhizosphere soil and plant roots are sampled and microbial communities determined for each of the three compartments can easily produce a contingency matrix [10, 11]. Based on the matrix, beta-diversity are usually calculated, showing the compositional similarity of microbial community between each other. Thereby, the distances between samples from bulk soil (B), rhizosphere soil (R) and plant endosphere (E) based on Bray-Curtis dissimilarity as the equation (1), where n_*ik*_ is the abundance of OTU *k* at the *i* and *j* samples.

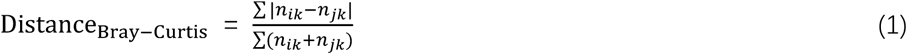

Based on the distances, a triangle can be constructed in a plane (Fig. 1). Consequently, the angle (α) size between the distance from bulk soil to endosphere (|BE|) and the distance from bulk soil to rhizosphere (|BR|) can be easily calculated from this ordinations.

**Fig. 1.**
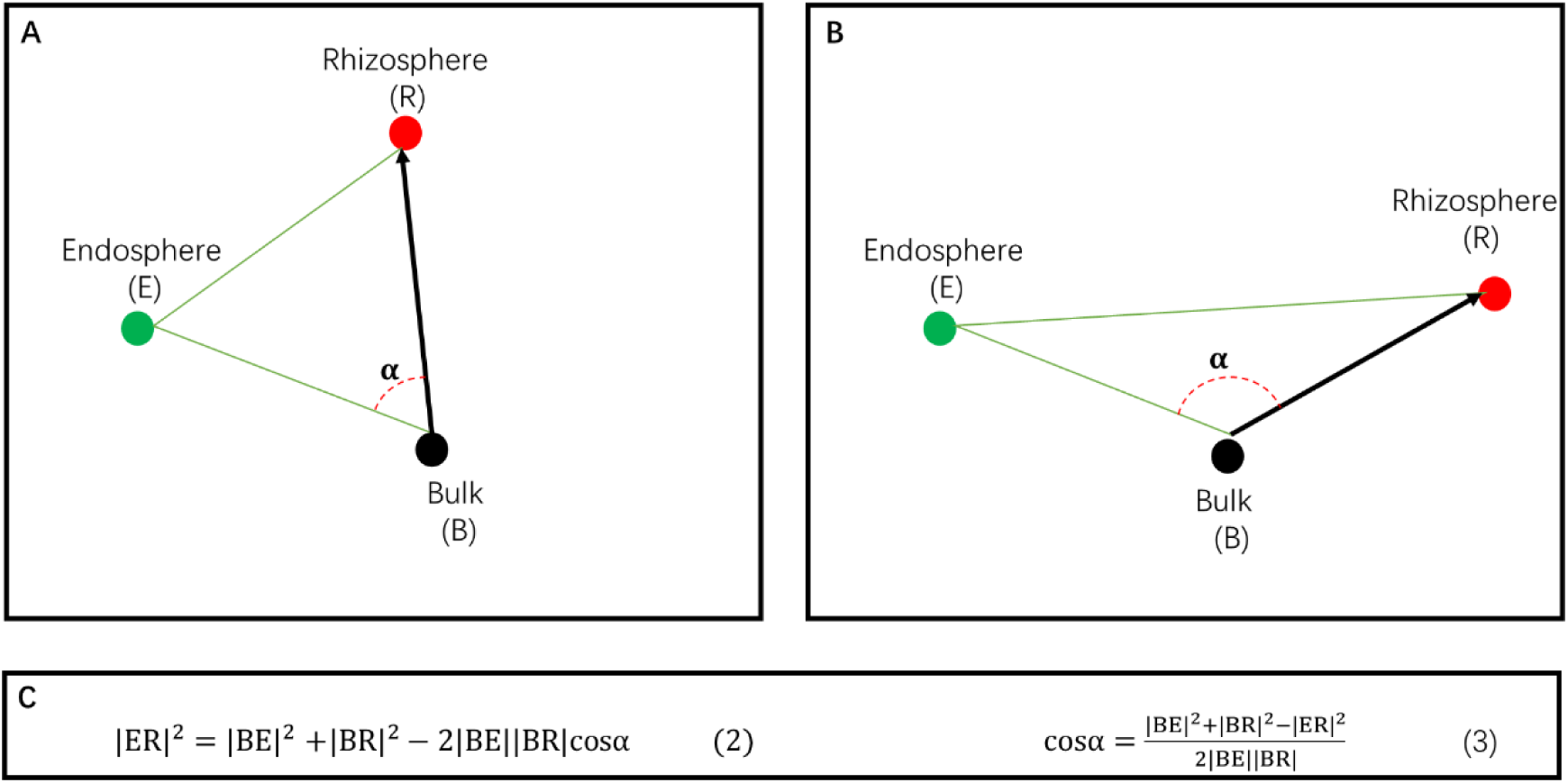
The triangle in a plane based on Bray-Curtis distances among the microbial communities’ structures in bulk soil (B), endoshphere (E) and rhizosphere (R). α is the angle degree between the |BE| and |BR|. Panels A and B indicate the two kinds of treatment arrangement in a plane. The back arrow indicates the assembling vector of rhizosphere microbiota recruiting from the bulk soil. Panels C shows the equations for calculating angle (α) size.

Then, we may link the endosphere (E) point with the bulk soil (B) point. Based on this line, an X-axis was constructed, then at the point B, a vertical line was drawn to be the Y-axis (Fig. 2). Thus, the vector representing the contribution of rhizosphere microbial assembly from the bulk soil (BR) can be decomposed along the X- and Y-axis. The resulting vector, along the X-axis, weights the effect from “plant selection”, since 1) the endophyte assemblages predominantly depends on the plant genotype, 2) the root endophyte assemblages overwhelmingly relies on the soil microbial reservoir since very few species stem from seed-borne or other negligible environments.

**Fig. 2.**
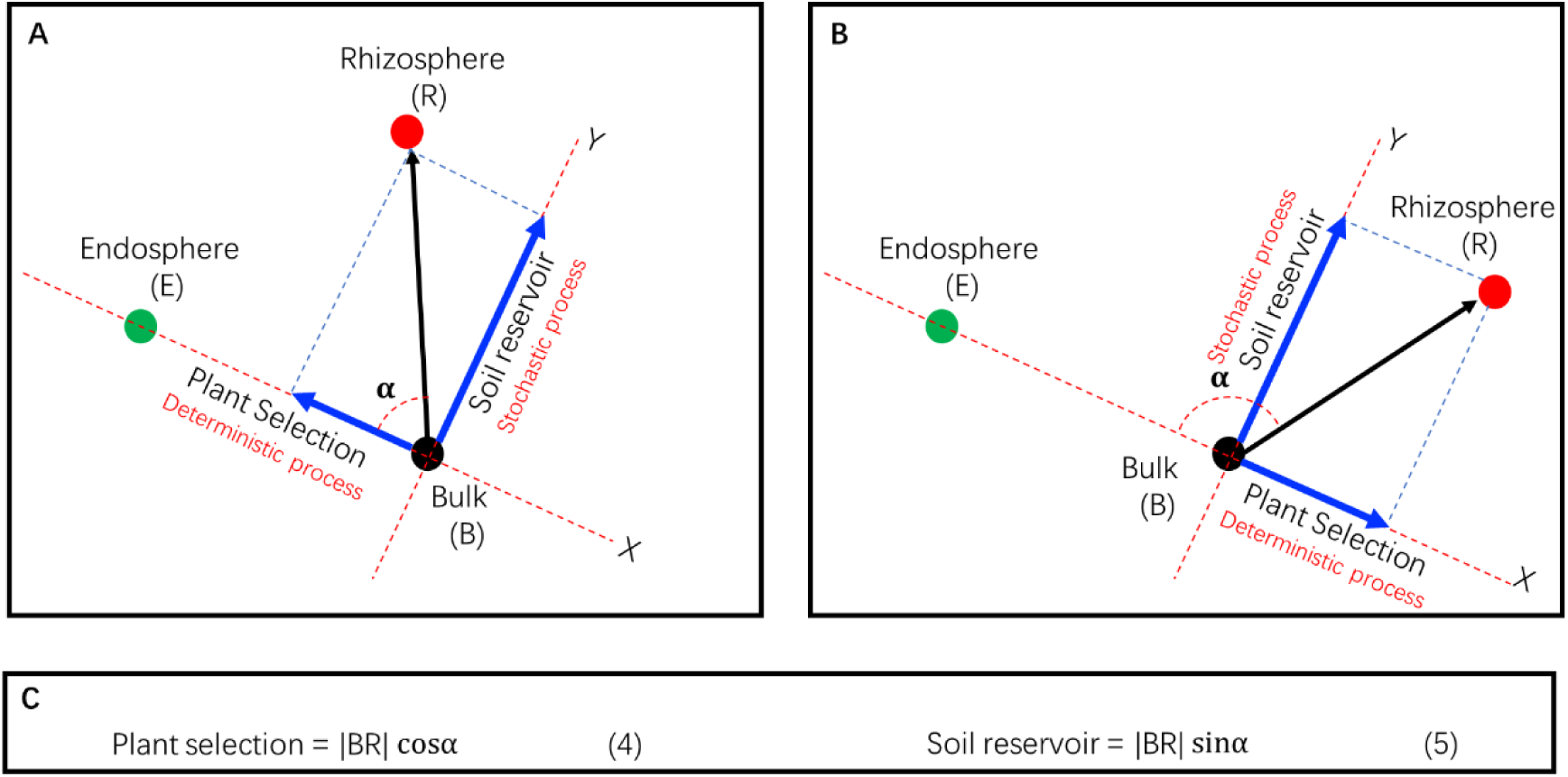
Decomposition of the rhizosphere microbial assembling forces into plant selection vector and Soil reservoir vector. X-axis was constructed based on the link line between bulk soil (B) and endosphere (E), and Y-axis was drawn as a vertical line at the point of B. the blue vectors indicate the partitioned vector of rhizosphere microbiota assembling. Panels A and B indicate the two kinds of treatment arrangement. Panels C shows the equations for “plant selection” and “soil reservoir”.

The other resulting vector, along the Y-axis and always pointing to the rhizosphere, indicates that the soil reservoir randomly influences microbial assembly in the rhizosphere. This stochastic process influenced by the soil microbial reservoir effect can be weighted as “Soil reservoir”.

As described above, we successfully weighted the major processes impacting the assembly of the rhizosphere microbiome due to plant selection (a deterministic process) and from the soil reservoir (a stochastic process). These effects were termed as “plant selection” and “soil reservoir”, respectively.

However, it is interesting to observe that: 1) if α < 90° (Fig.2A), the value of the plant selection effect can be positive, this means the vector points to the direction of the endosphere microbiome, indicating the plant selection confers a recruiting power for microbial assembly in the rhizosphere; 2) if α > 90° (Fig.2B), a negative value of plant selection is obtained, this means the vector points in the reverse direction in relation to the endosphere. This negative value indicates that plant selection is a repulsive effect in assemblage selection, perhaps due to incompatibility between plant growth and the indigenous microbiota. We suspect that this incompatibility may be observed in disease-conducive soils, as previous studies have reported the recruitment of beneficial microbes for fitness maintenance [12, 13]. Thus, if the bulk soil microbial reservoir is characterized by microbial taxa detrimental to plant fitness, a plant-derived negative selection may take place.

Overall, this proposed method provides a platform to partition contributions in rhizosphere microbiome assembly from plant selection (a deterministic process) and soil reservoir (a stochastic process). For example, this method can be used to quantify the soil reservoir effect with one cultivar plant in different soils or between different cultivars in one soil type. This weighting method may also provide a comparable parameter to evaluate soil health for plant growth.

## Acknowledgments

This work was supported by National Natural Science Foundation of China (31772398). We thank Dr. C. Ryan Penton of Arizona State University for the valuable comments.

## Author contributions

NL, PV and QRS designed the study. NL, CX, PV and QRS wrote the manuscript.

## R code accessibility

R code for |ER|, |BR|, |BE|, α, plant selection, and soil reservoir calculation based on Bray– Curtis dissimilarit can be provided by correspondence.

